# Voltage-based inhibitory synaptic plasticity: network regulation, diversity, and flexibility

**DOI:** 10.1101/2020.12.08.416263

**Authors:** Victor Pedrosa, Claudia Clopath

**Affiliations:** Department of Bioengineering, Imperial College London

**Keywords:** synaptic plasticity, inhibitory plasticity, spike-timing-dependent plasticity, voltage-based plasticity

## Abstract

Neural networks are highly heterogeneous while homeostatic mechanisms ensure that this heterogeneity is kept within a physiologically safe range. One of such homeostatic mechanisms, inhibitory synaptic plasticity, has been observed across different brain regions. Computationally, however, inhibitory synaptic plasticity models often lead to a strong suppression of neuronal diversity. Here, we propose a model of inhibitory synaptic plasticity in which synaptic updates depend on presynaptic spike arrival and postsynaptic membrane voltage. Our plasticity rule regulates the network activity by setting a target value for the postsynaptic membrane potential over a long timescale. In a feedforward network, we show that our voltage-dependent inhibitory synaptic plasticity (vISP) model regulates the excitatory/inhibitory ratio while allowing for a broad range of postsynaptic firing rates and thus network diversity. In a feedforward network in which excitatory and inhibitory neurons receive correlated input, our plasticity model allows for the development of co-tuned excitation and inhibition, in agreement with recordings in rat auditory cortex. In recurrent networks, our model supports memory formation and retrieval while allowing for the development of heterogeneous neuronal activity. Finally, we implement our vISP rule in a model of the hippocampal CA1 region whose pyramidal cell excitability differs across cells. This model accounts for the experimentally observed variability in pyramidal cell features such as the number of place fields, the fields sizes, and the portion of the environment covered by each cell. Importantly, our model supports a combination of sparse and dense coding in the hippocampus. Therefore, our voltage-dependent inhibitory plasticity model accounts for network homeostasis while allowing for diverse neuronal dynamics observed across brain regions.

## Introduction

Neuronal diversity has been observed across different brain regions^1–4^. In the rat hippocampus, this diversity is thought to be crucial to allow for the encoding of a behaviourally relevant range of environment sizes^1^. In the cortex, excitatory and inhibitory neuron firing rates are distributed across a wide range of possible values, with most neurons exhibiting a low firing rate and a few neurons firing at a very high firing rate^3^. This diversity in neuronal activity can emerge from heterogeneously connected excitatory and inhibitory neurons^5;6^.

Cortical neurons receive balanced excitatory and inhibitory inputs^7–10^. This balance between excitation and inhibition is thought to be important for network stability and signal processing^11–16^. Additionally, some cortical neurons have been shown to receive co-tuned excitatory and inhibitory inputs in a stimulus-specific manner^7;9;8;17^. The mechanisms that support and promote this balanced state in biological conditions, however, are still under intense debate. Inhibitory synaptic plasticity has been proposed as a potential candidate to fulfill this role^15;18–20^. By modulating inhibitory connections, the network can recover to a balanced state even in face of continuously-changing excitatory connections^15;21;22^. Any mechanism that modulates the balance between excitation and inhibition, however, should be able to support the neuronal diversity observed in biological neural networks.

A commonly used inhibitory synaptic plasticity model modulates inhibitory connections depending on the timing of pre- and postsynaptic spikes^15^. This spike-based inhibitory synaptic plasticity (sISP) rule regulates the balance between excitatory and inhibitory inputs while imposing a target firing rate for the postsynaptic neuron^15^. When combined with correlated excitatory and inhibitory inputs, this plasticity model produces co-tuned excitatory and inhibitory receptive fields^15;22^. Because of the restrictions that this model imposes onto postsynaptic firing rates, however, once the balanced state is achieved, responses to stimuli can only be perceived transiently^15^. The timescales at which these responses can be observed are determined by the timescales at which inhibitory synapses are updated. Moreover, in a recurrent network, the average firing rate of all excitatory cells converge to the same value, independently of their feedforward inputs.

Excitatory synaptic plasticity has been vastly explored and several plasticity models have been proposed, including spike-timing-dependent plasticity (STDP)^23^ and voltage-based models^24;25^. In contrast, inhibitory synaptic plasticity models have only recently started being investigated and the range of proposed plasticity models and applications is still limited^26;15;19;27^. Recent experimental data has suggested that the rules governing the change in inhibitory connections might depend on concurrent excitatory inputs^10^. Theoretical studies have shown that co-dependent excitatory and inhibitory synaptic plasticity rules can regulate network activity without setting a target firing rate for postsynaptic excitatory cells^28^. Moreover, accumulating evidence indicates that inhibitory plasticity rules are interneuron-type specific^29–31^, a characteristic that has been suggested to be important for controlling place field formation and consolidation in CA1 pyramidal cells^29^. Additionally, it has been reported experimentally that inhibitory synaptic plasticity in hippocampal CA1 interneuron synapses could only be induced by presynaptic theta burst stimulation if the postsynaptic membrane voltage was clamped at a hyperpolarized potential^29^. Under spike-timing-dependent protocols, inhibitory synaptic changes could not be observed using single postsynaptic spikes. Instead, inhibitory plasticity required postsynaptic bursts^29^.

We propose a voltage-based inhibitory synaptic plasticity (vISP) model in which the updates in inhibitory synaptic weights depend on the postsynaptic membrane voltage and presynaptic spikes. According to our plasticity model, inhibitory synaptic weights are updated to regulate the postsynaptic membrane voltage over a long timescale. We next explore the effects of this plasticity model using a feedforward network of excitatory and inhibitory inputs. We show that our model can modulate pyramidal cell activity by imposing a natural maximum firing rate. Contrary to previous models of sISP, however, our model does not impose a unique target postsynaptic firing rate. We then implement our model in a recurrent network and show that network heterogeneity can be observed in the neuronal activity. Finally, we introduce vISP in a model of the CA1 hippocampal region. Our model reproduces several experimental observations such as the co-tuning between excitation and inhibition in auditory cortex, the wide distribution of firing rates in hippocampal pyramidal cells, and the diverse range of place cell features. Therefore, our voltage-based model regulates network activity while allowing for a diversity in pyramidal cell firing rates.

## Results

### Voltage-based inhibitory synaptic plasticity model

In-vitro experiments suggest that inhibitory synaptic plasticity may depend on postsynaptic depolarization^29^. Moreover, current models of inhibitory plasticity impose strong homeostatic constraints in simulated neural networks^19^. We propose an inhibitory plasticity model in which the change in synaptic connections from inhibitory neurons onto pyramidal cells depends on the postsynaptic membrane voltage. In our voltage-based inhibitory synaptic plasticity model (vISP), the update in inhibitory weights aims to maintain the average postsynaptic membrane voltage at a target value over a long timescale (figure 1A, see methods). Additionally, to ensure a minimum level of neuronal activity, inhibitory presynaptic spikes lead to synaptic weight depression (iLTD).

**Figure 1:**
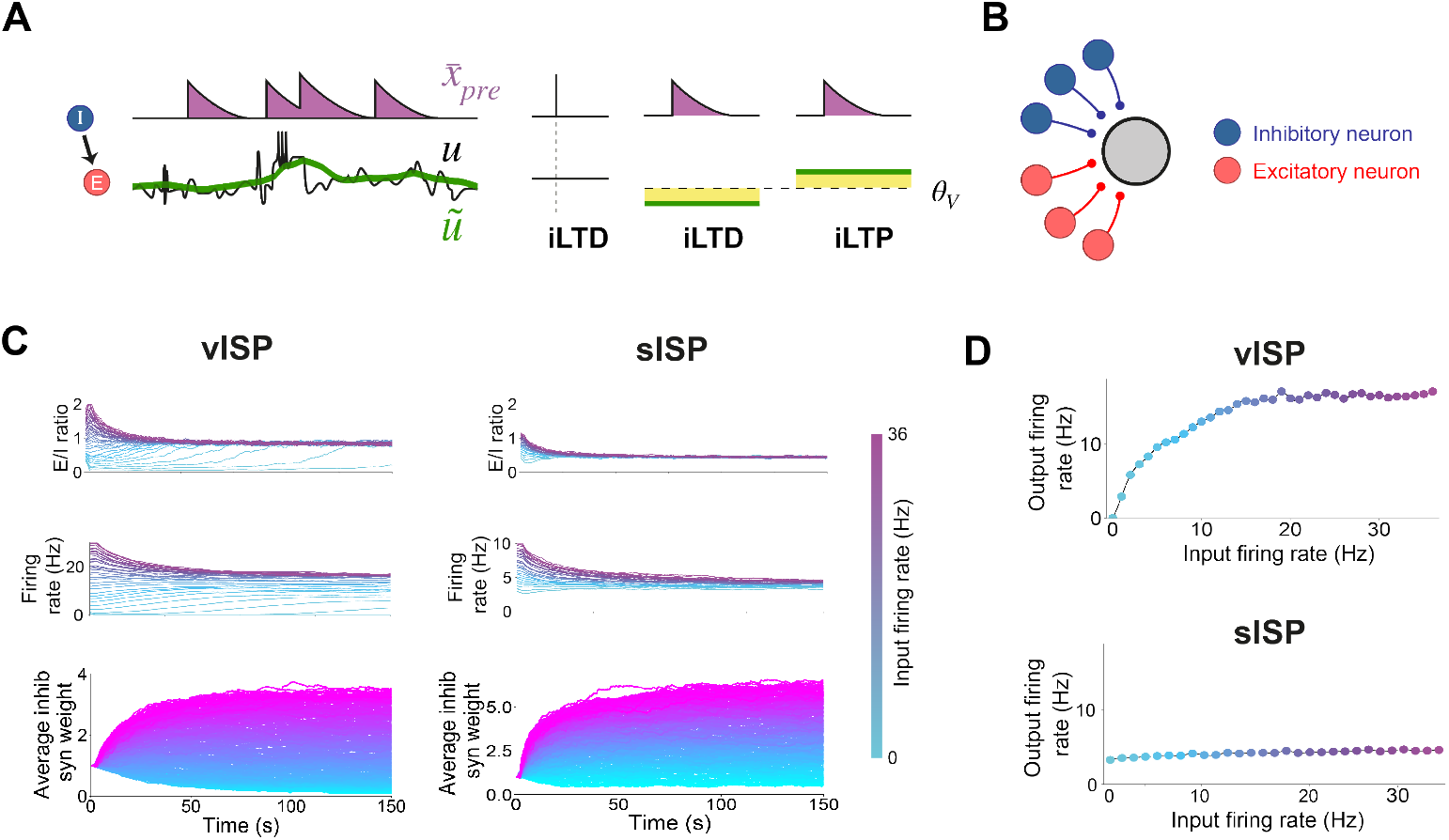
Voltage-dependent inhibitory synaptic plasticity regulates postsynaptic activity without setting a target firing rate. **(A)** Model diagram and variables. Our voltage-dependent inhibitory synaptic plasticity model (vISP) depends on a presynaptic trace, 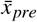, and a low pass-filtered version of the postsynaptic membrane voltage, 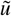. Presynaptic spikes lead to two independent processes: iLTD at the time of the spike, and either iLTD or iLTP during a short interval determined by the presynaptic trace 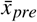. The polarity of this second term (i.e. iLTD or iLTP) depends on the value of 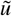 relative to a target value *θ_V_*. **(B)** Feedforward network diagram. We simulate a feedforward network composed of a set of excitatory (red) and inhibitory (blue) neurons projecting onto one postsynaptic excitatory cell (grey). The firing rates of input neurons vary over time but are kept, on average, at one specific value for excitatory neurons and at an independent value for inhibitory neurons. **(C)** Comparison between voltage-dependent inhibitory synaptic plasticity (vISP, left) and spike-based inhibitory synaptic plasticity (sISP, right). Top: E/I ratio (ratio between excitatory and inhibitory currents) as a function of time; middle: postsynaptic firing rate as a function of time; bottom: average inhibitory synaptic weight as a function of time. Each color corresponds to a simulation with one value of excitatory input firing rate. **(D)** Postsynaptic firing rate as a function of the mean firing rate of excitatory input neurons. The voltage-dependent inhibitory synaptic plasticity regulates postsynaptic firing rate by setting a maximum but not a target value. The spike-based inhibitory synaptic plasticity, however, imposes a unique target firing rate on the postsynaptic neuron regardless of the intensity of excitatory inputs.

### Voltage-dependent inhibitory synaptic plasticity regulates postsynaptic activity without setting a target firing rate

We first investigate whether our inhibitory plasticity model can regulate pyramidal cell firing rate. We simulate a feed-forward network composed of excitatory and inhibitory neurons projecting onto one postsynaptic neuron (figure 1B). The input neurons fire, on average, with the same firing rate and the synaptic weights from excitatory input neurons are all fixed at the same value. The weights for inhibitory synapses are initialized at a small value and are updated following the vISP model. We then perform these simulations for several levels of excitatory input firing rate. Independently of the initial conditions, the vISP model regulates the inhibitory connections such that the ratio between excitatory and inhibitory currents onto the postsynaptic cell (E/I ratio) converges to the same level (figure 1C). Interestingly, our inhibitory plasticity rule sets a natural maximum firing rate for the postsynaptic neuron (figure 1C-D). Importantly, this model does not restrict the postsynaptic activity to a narrow range. Instead, the postsynaptic firing rate can assume any value from zero to the maximum firing rate imposed by the inhibitory plasticity rule (figure 1D). Therefore, the vISP model regulates pyramidal cell activity by setting a target E/I ratio and a maximum firing rate without over-constricting the postsynaptic activity.

To compare the effects of our model with previous models, we perform the same simulations replacing our inhibitory plasticity model with a spike-based inhibitory plasticity rule^15^. Under this rule, near-coincident pre- and postsynaptic spikes lead to synaptic potentiation whereas each presynaptic spike leads to synaptic depression (see methods). This synaptic plasticity rule has been shown to set a target firing rate for the postsynaptic neuron^15^. Indeed, in our simulations, the sISP model sets a target value for both the E/I ratio and the postsynaptic firing rate (figure 1C-D). This difference from the vISP rule is observed even though the evolution of the average inhibitory synaptic weight does not differ, qualitatively, from the one observed in our model. Therefore, the sISP model regulates pyramidal cell activity by constraining the postsynaptic firing rate to a narrow range around the target firing rate.

### vISP and correlated E-I inputs lead to co-tuned excitatory and inhibitory receptive fields

We next investigate the effect of the vISP model on inhibitory receptive field formation. In particular, we wonder whether this inhibitory plasticity model can account for the co-tuning of excitatory and inhibitory currents observed in cortical neurons^7;9;17;8;14^. To address that, we simulate a feedforward network of excitatory and inhibitory neurons projecting onto one postsynaptic neuron. Those two populations of neurons are organized in pairs such that each pair of excitatory and inhibitory neurons fire with the same time-varying firing rate (figure 2A). This is equivalent to simulating pairs of excitatory and inhibitory neurons receiving similar inputs. The excitatory synaptic weights are initialized such that the excitatory receptive field is Gaussian shaped with a peak at input neuron 10. Those synaptic weights are kept fixed throughout the simulations. The inhibitory synaptic weights are initialized at a low value and evolve following the vISP model. Similarly to the case with homogeneous excitatory connections, the change in inhibitory connections regulates the E/I ratio, forcing it towards a target value close to 1 (figure 2C) and the post-synaptic firing rate stabilizes at a low level (figure 2D). Interestingly, although the vISP model allows for a wide range of postsynaptic firing rates, the average inhibitory currents converge to the same values of their corresponding excitatory counterparts (figure 2B and 2E). Therefore, the vISP model supports the emergence of a co-tuning between excitation and inhibition.

**Figure 2:**
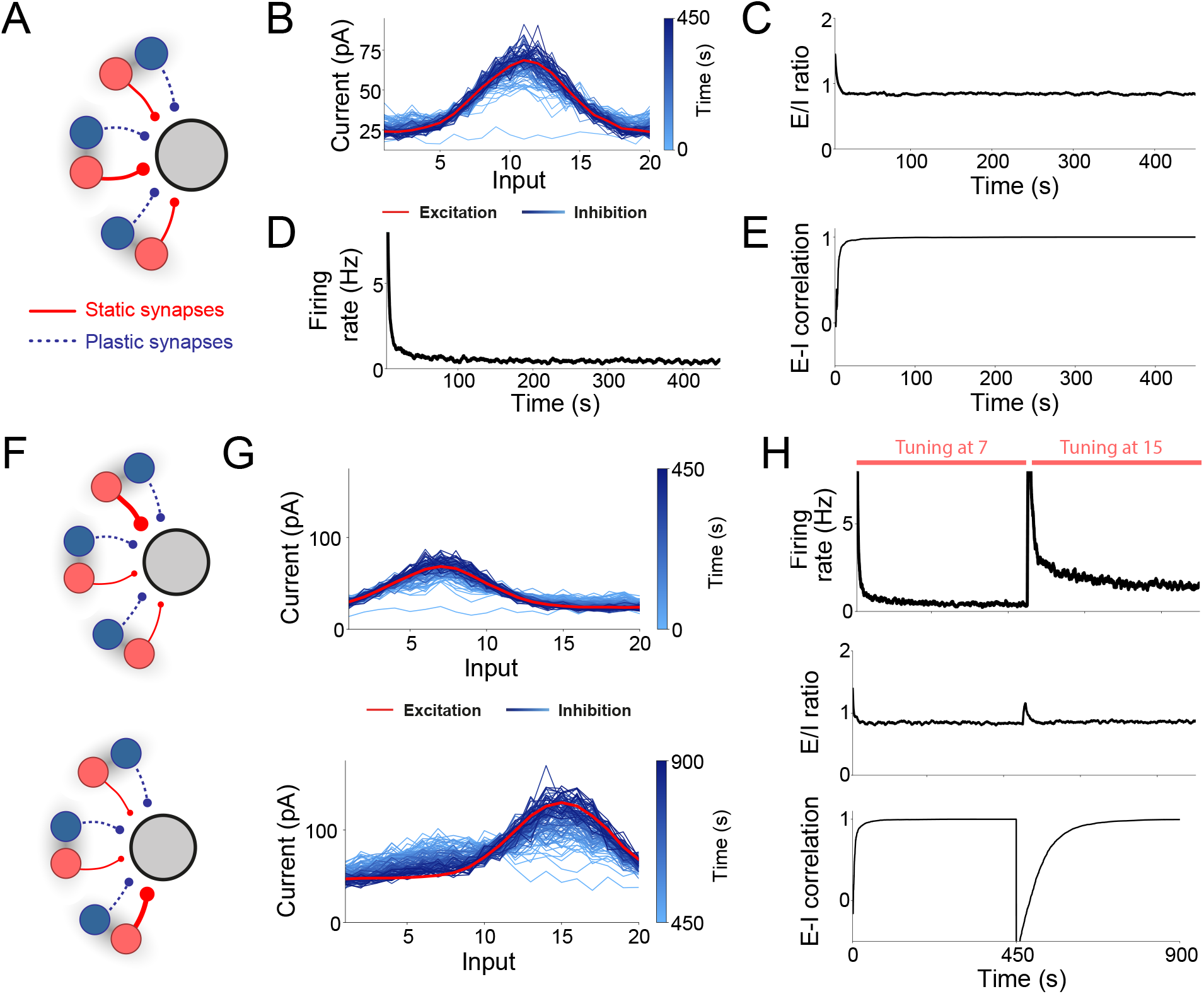
vISP and correlated E-I inputs lead to co-tuned excitatory and inhibitory receptive fields. **(A)** Network diagram. We simulate a feedforward network composed of a set of excitatory (red) and inhibitory (blue) neurons projecting onto one postsynaptic excitatory cell (grey). Each excitatory neuron is associated with an inhibitory counterpart such that they follow the same time-varying firing rate (grey shading). The firing rates of different pairs of input neurons vary over time but are kept, on average, at the same level. Inhibitory connections follow our voltage-based inhibitory synaptic plasticity (vISP) model. **(B)** Evolution of inhibitory receptive field. Synaptic weights for each inhibitory (blue) and excitatory (red) input neuron. The different shades of blue represent snapshots of the inhibitory weights at different times throughout the simulation. **(C)** Ratio between excitatory and inhibitory currents onto the postsynaptic cell as a function of time. The curve is smoothed using a rolling average over 100 ms. **(D)** Postsynaptic firing rate as a function of time. **(E)** Correlation between excitatory and inhibitory receptive fields as a function of time. **(F-H)** In these simulations, the excitatory receptive fields (synaptic weights) are kept constant for half of the simulation (first 450 s). The synaptic weights are then abruptly modified such that the strongest synaptic weight is moved from input 7 to input 15 and the overall synaptic weight amplitude is twice as strong. The excitatory synaptic weights are then kept constant for the second half of the simulation (last 450 s). **(F)** Network diagram for the first (top) and second (bottom) half of the simulation. **(G)** Evolution of inhibitory synaptic weights following a sudden change in excitatory receptive field. Synaptic weights for each inhibitory (blue) and excitatory (red) input neuron for the first (top) and second (bottom) half of the simulation. The different shades of blue represent snapshots of the inhibitory weights at different times throughout the simulation. **(H)** Postsynaptic firing rate (top), E/I ratio (middle), and E-I correlation (bottom). Following the sudden change in excitatory synaptic weights, the postsynaptic firing rate transiently increases and slowly returns to a lower level. Importantly, the final postsynaptic firing rate is higher than in the first half of the simulation. E-I correlation is measured as the correlation between excitatory and inhibitory receptive fields (synaptic weights) as a function of the input. E/I ratio and E-I correlation converge converge to the same level, regardless of the amplitude of excitatory inputs.

The co-tuning between excitatory and inhibitory receptive fields has also been observed using a spike-timing dependent inhibitory plasticity ruleor its rate-based version. To test this under the same conditions as the ones used with the vISP model, we perform the same simulations as before while replacing the voltage-dependent inhibitory plasticity rule with a spike-based model. As expected, the inhibitory synaptic inputs converge to the same levels as their corresponding excitatory input (supp. figure 1). In summary, both voltage- and spike-based inhibitory plasticity rule account for the development of co-tuned excitatory and inhibitory receptive fields when excitatory and inhibitory inputs are correlated.

### Inhibitory connections adapt to changes in excitatory input while allowing for diversity in pyramidal cell firing rate

As excitatory synaptic weights are constantly changing^32–39^, neural circuits should be able to adapt to these changes. In particular, sensory stimulation paired with the activation of neuromodulatory circuits has been shown to induce changes in excitatory receptive fields^9;40^. This change is then followed by an adaptation of inhibitory receptive fields in order to restore the co-tuning between excitation and inhibition observed in auditory cortex^9;40^. Importantly, the firing rate following the restoration of balance does not necessarily correspond to the level before induction of plasticity^40^. To test whether a network governed by the vISP model can adapt to changes in excitatory synaptic weights, we simulate a feed-forward network analogous to the one we simulated before but with an excitatory receptive field centered around input 7 (figure 2F). The activities of excitatory and inhibitory neurons associated to the same input index are correlated in time. The excitatory synaptic weights are kept fixed while the inhibitory synaptic weights evolve following the vISP model. Similarly to what we observed in the previous case, the inhibitory receptive field converges to match the excitatory inputs (figure 2G). Consequently, the postsynaptic firing rate decays to an almost-silent stage (figure 2H). After 450 s, the excitatory synaptic weights are shifted instantly such that the peak of the excitatory receptive field becomes centered around input 15 and its amplitude is twice the original amplitude (figure 2F). This simulates the natural change in excitatory tuning observed in auditory cortex following neuromodulatory stimulation^9;40^. The excitatory synaptic weights are then kept constant for another 450 s while the inhibitory synaptic weights are allowed to change. The inhibitory receptive field adapts to the changes in excitatory inputs moves towards the new excitatory receptive field (figure 2G). Remarkably, the postsynaptic firing rate does not return to the same level. Instead, it decays to a higher level when compared to the first 450 seconds of simulation (figure 2H). Meanwhile, the ratio between excitatory and inhibitory currents and the correlation between excitatory and inhibitory receptive fields return to baseline level (figure 2H). Contrastingly, although the sISP model supports the co-tuning between excitation and inhibition, it constrains the postsynaptic firing rate to a narrow range around a target value (suppl fig 2F-H). Therefore, the vISP model provides a mechanism with which a network can adapt to changes in excitatory inputs and still allow for diversity in neuronal activity.

### Memory formation and retrieval is supported by vISP

Our voltage-based-inhibitory synaptic plasticity does not set a target firing rate in a feedforward network. To test whether this can be extended to a recurrent network and its functional consequences, we simulate a randomly connected network of excitatory and inhibitory neurons. All the excitatory connections are kept fixed throughout the simulations whereas inhibitory connections onto excitatory cells follow our vISP model (figure 3A). We then select a subset of the excitatory cells and increase the connections amongst them to simulate memory formation by instantly multiplying synaptic weights by 3. After network stabilization, we test memory retrieval by stimulating half of the neuronal assembly and recording the activity of the other half of the assembly.

**Figure 3:**
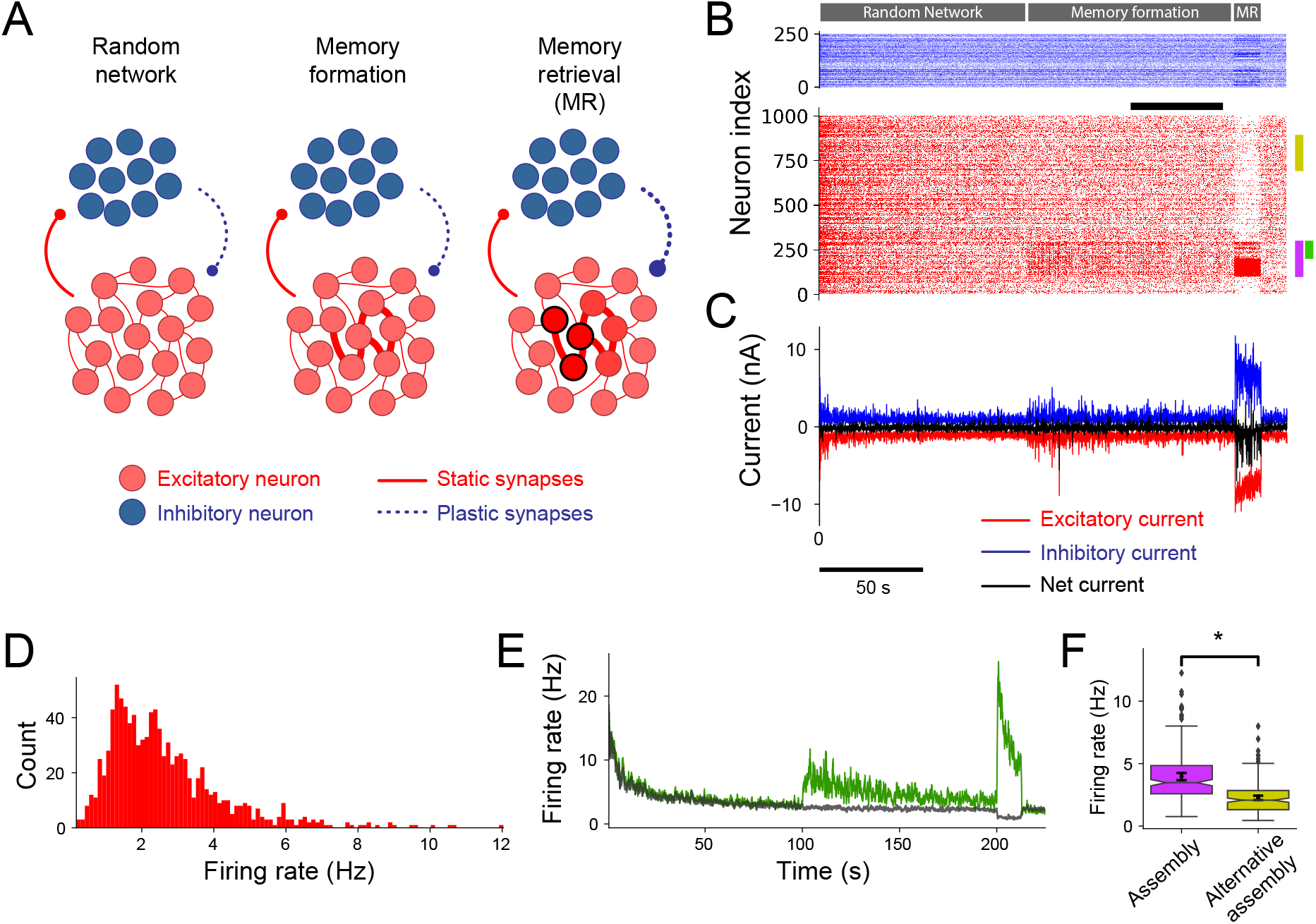
Memory formation and retrieval is supported by vISP. **(A)** Network diagram and simulation protocol. We simulate a recurrent network of excitatory and inhibitory neurons. All the neurons are randomly connected and receive inputs from a set of input neurons randomly connected to the neural network. Excitatory connections are static whereas inhibitory connections onto excitatory cells are plastic and follow our voltage-dependent inhibitory synaptic plasticity rule. The network is first simulated and reaches a stable state (left). A subset of excitatory neurons are then selected and the existing connections amongst these neurons are strengthened, multiplied by 3 (middle, memory formation). Finally, after the network reaches a new stable state, half of the neurons selected in the previous stage are stimulated by an extra set of input neurons (right, memory retrieval). Inhibitory connections are plastic throughout the entire simulation. **(B)** Inhibitory (blue) and excitatory (red) spikes for one example simulation. The memory formation stage starts at 100 seconds and the extra input (memory retrieval) is applied at 200 seconds for 5 seconds. Purple: memory assembly whose neurons are strongly connected amongst themselves after *t* = 100 s. Green: half of the neurons in the memory assembly that does not receive an extra external input at *t* = 200 s. Yellow: an alternative set of cells of the same size as the memory assembly. The grey bar indicates the interval over which the average firing rates are measured in D and F. **(C)** Inhibitory (blue), excitatory (red), and net (black) currents received by a random neuron within the second half of the memory assembly (green bar in B). Please note that we use the convention that inhibitory currents assume positive values. **(D)** Distribution of firing rates measured between 150 and 190 seconds (grey bar in B) across all excitatory neurons. **(E)** Firing rate for the second half of the neurons in the memory assembly (green bar in B) and background activity for the neurons outside of the memory assembly. **(F)** Comparison between the average firing rate of the memory assembly (purple bar in B) and an alternative set of cells (yellow bar in B). The average firing rate is measured from 150 to 190 seconds. The firing rates are significantly different even after network stabilization (p <10^−4^, Kruskal-Wallis H-test, *n* = 200 neurons).

Since all the neurons are randomly connected and receive random inputs, the network activity is homogeneous during the first stage of the simulation (figure 3B and suppl figure 2A). The inhibitory synaptic connections then increase and bring the activity of the entire network to a low level (figure 3B and suppl figure 2A). Once the connections between pairs of neurons in the assembly are increased, the excitatory and inhibitory currents received by a neuron in the assembly increase (figure 3C). The increase in excitatory current is higher, leading to an increase in the firing rate of neurons in the memory assembly (figure 3B,E). The inhibitory connections then undergo plasticity, lowering the firing rate of the memory assembly (figure 3B,E). Notably, the firing rate of the assembly does not return to the same level of the rest of the network (figure 3B, E, and F and suppl figure 2A) and the firing rates of the neurons in the excitatory network are widely distributed (figure 3D). In contrast, when the same simulations are performed using a spike-based inhibitory synaptic plasticity model, the activity of the neuronal assembly is indistinguishable from the activity of the rest of the network at the steady-state (suppl figure 3). Although vISP does not force the firing rate of the memory assembly to return to baseline level, the average ratio between excitatory and inhibitory currents converges to a unique level (suppl figure 2C). Therefore, our vISP model regulates the activity of memory assemblies but does not force it to fade completely to baseline level.

We next investigate whether we can retrieve the memory using external stimulation. We increase the external input to half of the neurons forming the memory assembly and record the activity of the remaining neurons within the assembly. Similarly to what is observed with a sISP model (suppl figure 3A-B), the partial memory stimulation leads to an increase in the activity of the remaining neurons in the assembly (figure 3B,E). This increase is also accompanied by a decrease in the activity of the excitatory neurons outside of the memory assembly (figure 3B,E). This decrease, however, may depend on the level of lateral inhibition in the network. In summary, vISP supports memory formation and retrieval via pattern completion. Moreover, neurons that are part of a memory assembly (i.e. neurons that are strongly connected) show a higher firing rate even after network stabilization.

### vISP supports diversity in a heterogeneous recurrent network

To extend our investigation into the functional consequences of our vISP model in recurrent networks, we next simulate a heterogeneously connected neural network. In these simulations, each excitatory neuron has a propensity of forming connections with other excitatory neurons whereas all the other synaptic connections are randomly assigned (figure 4A-B). Once assigned, all the excitatory connections are kept constant throughout the entire simulation whereas inhibitory synapses follow our vISP model. During the first stages of the simulation, the highly connected neurons fire at a much higher firing rate when compared to the overall network activity (figure 4C,F). The changes in inhibitory synaptic weights due to vISP then push these neurons to a lower firing rate (figure 4F). Different to what is observed in simulations using a sISP model (suppl figure 4E), however, the final firing rates are broadly distributed, ranging from zero to above 10 Hz (figure 4D). The more the propensity of a neuron being connected to other neurons, the higher its firing rate (figure 4E). Therefore, although the vISP model regulates the overall network activity, it supports a diversity in neuronal firing rates.

**Figure 4:**
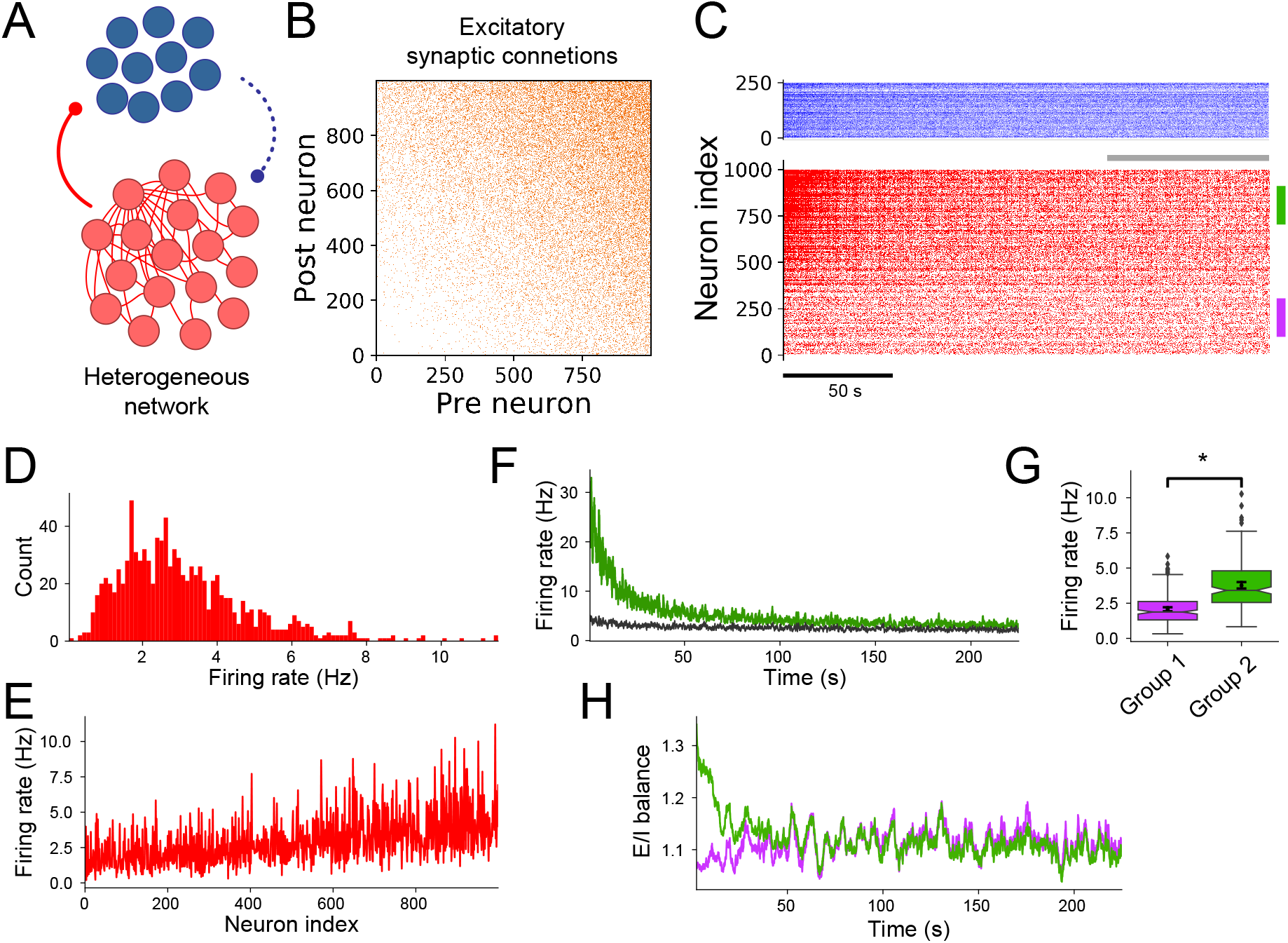
vISP supports diversity in a heterogeneous recurrent network. **(A)** Network diagram. Excitatory neurons are heterogeneously connected to other excitatory neurons and randomly connected to inhibitory neurons. Inhibitory neurons are also randomly connected to other inhibitory neurons and to excitatory neurons. All the excitatory connections are kept fixed whereas inhibitory connections onto excitatory cells are plastic and follow our voltage-dependent inhibitory synaptic plasticity rule. **(B)** Excitatory synaptic connectivity matrix. All the connections have the same strength, but the neurons are not uniformly connected. Neurons are sorted by their propensity of forming connections with other neurons. **(C)** Inhibitory (blue) and excitatory (red) spikes. Purple: set of 200 neurons with low propensity of forming connections with other neurons in the network. Green: set of 200 neurons with high propensity of forming connections with other neurons in the network. The grey bar indicates the interval over which the average firing rates are measured in D, E and G. **(D)** Distribution of firing rates measured from 150 and 220 seconds (grey bar in C) across all excitatory neurons. **(E)** Firing rates shown in D for each neuron. Neurons are sorted by their propensity of forming connections with other neurons. **(F)** Firing rate for a set of highly connected neurons (green bar in C) and background activity measured across all the other neurons. **(G)** Comparison between the average firing rate of a set of weakly connected neurons (group 1, purple bar in C) and a set of highly connected neurons (group 2, green bar in C). The average firing rate is measured from 150 to 220 seconds. The firing rates are significantly different even after network stabilization (*p* < 10^−4^, Kruskal-Wallis H-test, *n* = 200 neurons). **(H)** Average ratio between excitatory and inhibitory currents for a set of highly connected (green) and weakly connected (purple) neurons. The averages are measured across the 200 neurons in each group.

To analyze this diversity in more detail, we consider two subgroups of the network of excitatory neurons. The first group (group 1) is composed of neurons with a low propensity of forming connections with other neurons (figure 4C, purple bar). The second group (group 2) is composed of neurons with a high propensity of forming connections (figure 4C, green bar). After the simulation reaches a steady state, we measure the average firing rates of the neurons in each of these groups. The final firing rates of neurons in group 1 are significantly lower than the firing rates of the neurons in group 2 (figure 4G) whereas the average ratio between excitatory and inhibitory currents converge to the same level for both groups (figure 4H). If the inhibitory connections follow, instead, a sISP model, the firing rates of both groups converge to the same value (suppl figure 4C-D). Therefore, highly connected neurons are more active than weakly connected neurons if inhibitory connections follow a voltage-dependent inhibitory synaptic plasticity model. In summary, network diversity is preserved in a heterogeneous network following our vISP model.

### Heterogeneous excitability and vISP account for CA1 place field diversity

Heterogeneous neural networks have been observed across several brain regions^3;1;2^. One region in which this heterogeneity is particularly striking is the hippocampal CA1 sub-region. When an animal explores an environment, a subset of pyramidal cells remains silent while another subset becomes place cells. Moreover, amongst the place cells, neurons can show a plethora of behaviours in terms of overall firing rate, number of place fields, proportion of the environment covered by each cell, and the sizes of place fields^2;41^. This variability in behaviour has been associated with differences in intrinsic neuronal properties in rats^42^. Here, we investigate whether our vISP model supports the development of heterogeneous place fields while enforcing network homeostasis.

First, we simulate a network of CA1 pyramidal cells receiving lateral inhibition (figure 5A). Given that the connection probability between CA1 pyramidal cells is extremely low, we assume that there are no direct connections between excitatory CA1 cells. Connections from excitatory to inhibitory neurons and vice-versa are assigned randomly and their amplitudes are drawn from a log-normal distribution. While all the excitatory connections are kept fixed, inhibitory connections follow our vISP model. CA1 pyramidal cells also receive feedforward input from CA3 pyramidal cells, whose activity is spatially tuned. Each CA3 cell has a unique place field and the place fields of all the input neurons span the entire linear track (see methods). Finally, to model neuronal diversity, we assume that each excitatory neuron has a specific neuronal excitability. To that end, we assign to each CA1 pyramidal cell a fixed spiking threshold drawn from a uniform distribution. This excitability is kept constant throughout the simulations but varies from neuron to neuron (figure 5A).

**Figure 5:**
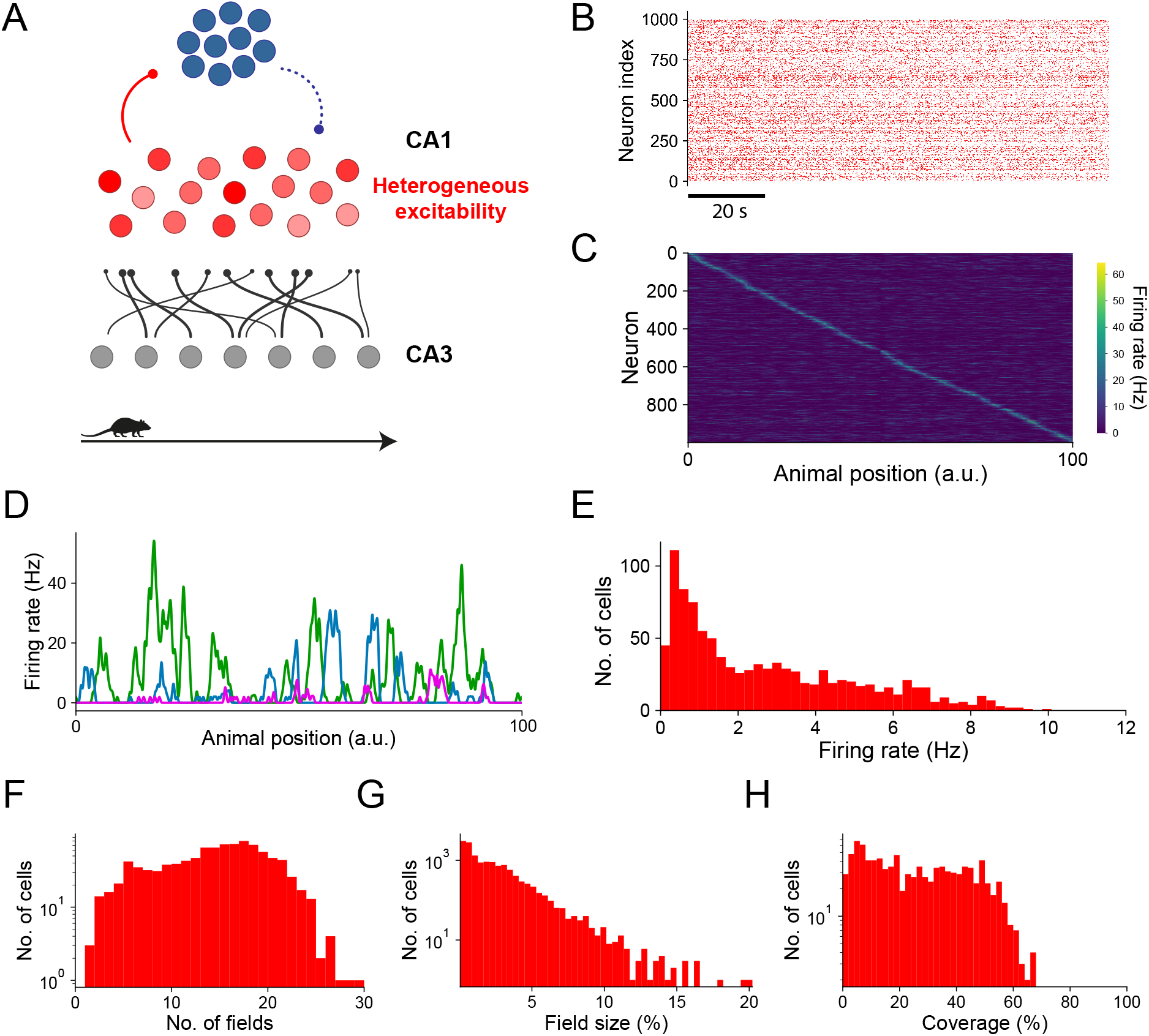
Heterogeneous excitability and vISP account for CA1 place field diversity. **(A)** Network diagram. CA1 pyramidal cells (red) receive inputs from CA3 excitatory cells (grey) and are recurrently connected via inhibition (blue). Each excitatory cell is assigned a level of excitability such that their spiking thresholds are drawn from a uniform distribution. CA3 place fields are uniformly distributed and spam the entire linear track. Connections from CA3 to CA1 pyramidal neurons and from pyramidal neurons to interneurons are random and fixed. The inhibitory connections onto CA1 pyramidal cells are initialised randomly and follow our voltage-dependent inhibitory plasticity rule. **(B)** CA1 pyramidal cell spikes for the entire simulation. The animal travels across the entire linear track in 10 s and is placed back at the initial position instantly. **(C)** CA1 pyramidal cell activity as a function of the animal position. Neurons were sorted by the position of their maximum firing rate. The activity is an average over the final 5 laps of simulation. CA1 place fields span over the entire linear track. **(D)** Example place fields for 3 CA1 pyramidal cells (each color represents one cell). Each cell can have multiple place fields across the track and their amplitudes can vary significantly. **(E)** Distribution of average firing rates across CA1 pyramidal cells. Firing rates were measured over the final 5 laps of simulation. The vISP model allows for a wide range of neuronal activities. **(F)** Distribution of number of fields. **(G)** Distribution of field size compared to the total track. **(H)** Distribution of the total coverage of each neuron compared to the total track. The vISP model combined with heterogeneous CA1 pyramidal cell excitability accounts for place cell diversity.

Similarly to what we observed in our previous simulations, inhibitory connections undergo plasticity, leading to a modulation of the overall network activity (figure 5B). Following stabilization, CA1 pyramidal cells develop reliable place fields that span the entire environment (figure 5C). Each cell, however, can have multiple place fields and the amplitudes of these fields can vary drastically from field to field (figure 5D). Interestingly, the average neuronal firing rate across the entire linear track can take a wide range of values (figure 5E), in agreement with experimental observations^1;2^. A small proportion of cells are completely silent while the animal traverses the track whereas other cells can have an average firing rate of up to almost 12 Hz (figure 5E). Importantly, this wide distribution of firing rates reflects into a wide range of place fields per cell (figure 5F), field sizes (figure 5G), and the neuron’s coverage of the track (figure 5H), in agreement with experiments^2;41^. Contrarily, a heterogeneous CA1 network in which inhibitory synapses follow a sISP model exhibit a narrow range of overall neuronal firing rates (suppl figure 5E). Consequently, the heterogeneity of this neural network is not manifested in the number of place fields per cell (suppl figure 5F) or how much of the track is covered by each cell (suppl figure 5H). Therefore, our voltage-dependent inhibitory synaptic plasticity model, combined with heterogeneous intrinsic CA1 pyramidal cell excitability, accounts for the experimentally-observed variety of place fields.

### vISP and heterogeneous excitability predict CA1 neuron propensity to develop place fields while supporting remapping

The most important characteristic of our vISP model is not only to allow for network diversity but, arguably, to allow for adaptation following network changes. To explore this aspect of our model, we simulate a hippocampal network while an animal changes from environment A to environment B (figure 6A). The simulated network is identical to the network simulated in the previous section. To simulate the environment change, we redraw the feedforward connections from CA3 to CA1 pyramidal cells from the same distribution as in environment A (figure 6A). This leads to a reshuffle of CA1 place fields (figure 6B) even though individual place fields and firing rate distributions are very similar to the ones measured in environment A (figure 6C-D).

**Figure 6:**
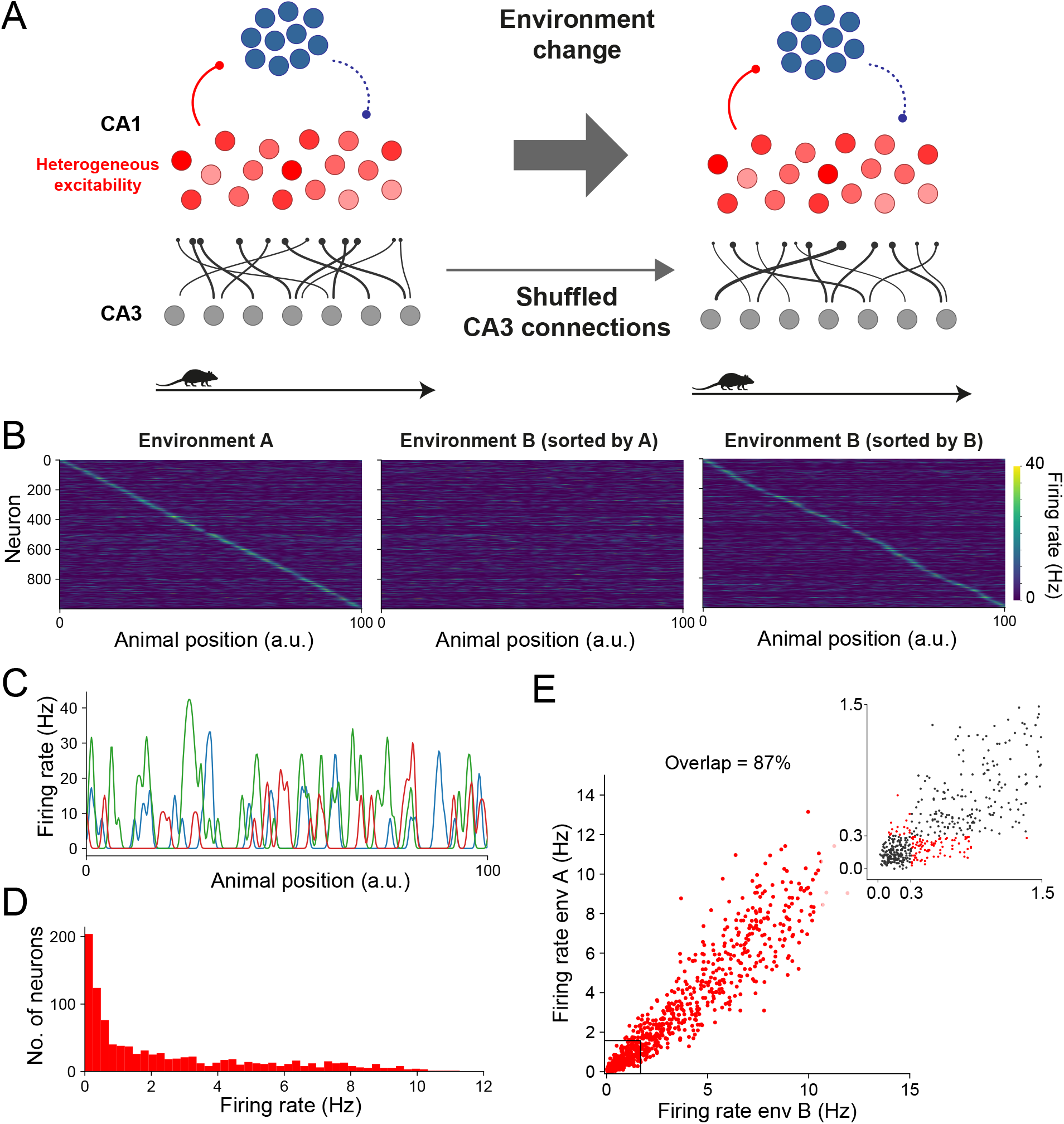
vISP and heterogeneous excitability predict CA1 neuron propensity to develop place fields while supporting remapping. **(A)** Network diagram and simulation protocol. The network is simulated as in figure 5. After half of the simulation time, the connections from CA3 to CA1 pyramidal cells are instantly re-drawn from the same distribution as before to simulate an environment change. The network is then simulated as in the first half of the simulation. All the other parameters are kept constant throughout the entire simulation, including the excitability of CA1 pyramidal cells. **(B)** CA1 pyramidal cell activity as a function of the animal position for the first half (environment A) and the second half (environment B) of the simulation. Neurons were sorted by the position of their peak in either environment A (left and middle panels) or environment B (right panel). The activity shown is an average over the final 5 laps of simulation in each environment. **(C)** Example place fields for 3 CA1 pyramidal cells in environment B (each color represents one cell). Similarly to before the environment change, each cell can have multiple place fields across the track and their amplitudes can vary significantly. **(D)** Distribution of average firing rates across CA1 pyramidal cells in environment B. Firing rates were measured over the final 5 laps of simulation. **(E)** Firing rate in environment A versus firing rate in environment B for all the CA1 pyramidal cells. The overlap represents the proportion of cells that are either active or silent in both environments (i.e. the proportion of cells that do not change their status as silent or place cells). Therefore, 13% of the pyramidal cells are only active in one of the environments. Inset: firing rate within the range between zero and 1.5 Hz. Red dots in the inset show the neurons that are active in only one of the environments. Neurons with an average firing rates below 0.3 Hz were considered silent.

We next investigate whether the change in environment (i.e. the change in CA3 to CA1 connections) leads to a change in the cells encoding the track. To that end, we compare the average activity of each cell in environment A with its activity in environment B (figure 6E). Most cells are active in both environments and their activities in environment B are correlated with their activities in the first environment, in agreement with experiments^1^. A small number of cells (13% of the CA1 pyramidal cells), however, are active in only one of the environments (figure 6E). This set of cells, albeit small in our simulations, could provide a mechanism for quick environment identification. In contrast, in a hippocampal network in which inhibitory synapses are governed by sISP, pyramidal cells are either active or silent in both environments (figure 6E). This network, therefore, might lack a mechanism for easy environment or context detection, limiting its consistency with experimental observations. Thus, our inhibitory plasticity model and neuronal excitability can account for the diversity in the propensity of neurons in developing place cells while allowing for CA1 pyramidal cell remapping.

## Discussion

We propose a voltage-based inhibitory synaptic plasticity model as a mechanism to regulate network activity. Our model imposes a target value for a low-passed-filtered version of the postsynaptic membrane potential. This allows for short-term fluctuations while imposing a long-term regulation of the neuron’s membrane potential. We analyse the effect of our inhibitory plasticity model on a feedforward network composed of excitatory and inhibitory neurons projecting onto one single neuron. Our vISP model regulates the postsynaptic activity by imposing a natural maximum firing rate and a unique target value for the ratio between excitatory and inhibitory inputs. Surprisingly, the vISP model does not over-restrict the postsynaptic firing rate. Instead, the postsynaptic neuron firing rate is only bounded by a maximum firing rate but can assume any value between zero and this maximum bound. This result contrasts with previous results observed under spike-based inhibitory synaptic plasticity rules^15;21;19;18^. When applied to a feed-forward network with correlated inhibitory and excitatory inputs, our vISP model leads to co-tuned excitatory and inhibitory receptive fields, as observed in auditory cortex^9;40^. Additionally, this co-tuning could be combined with a flexibility in the postsynaptic firing rate following changes in the excitatory receptive field. This feature might be essential when excitatory receptive fields change both tuning and amplitude, as observed in auditory cortex neurons following noradrenergic release^40^

In recurrent neural networks, inhibitory synaptic plasticity has been shown to be a good candidate to regulate network activity while supporting the existence of multiple cell assemblies^15^. When the connections between clusters of neurons—the cell assembly—are strengthened to store a memory, the sISP model leads to the potentiation of inhibitory connection onto the cell assembly. This strengthening of inhibitory connections ultimately pushes the cell assembly activity back to baseline level^15^. At this stage, the memory is stored in the synaptic weights but the activity of the cell assembly is indistinguishable from the activity of the rest of the network. Contrarily, if inhibitory connections follow our vISP model, the activity of the cell assembly does not return to baseline level. The activity of the memory assembly can, therefore, be differentiated from the activity of the other neurons in the network. For both plasticity models, partial stimulation of the memory assembly leads to pattern completion and the reactivation of the remaining neurons in the assembly. Notably, the level of activity in the assembly is a direct consequence of the amount of excitatory drive received by those neurons. The activity of these neurons might be indistinguishable from the activity of other neurons if all neurons are part of the same number of memory assemblies. Furthermore, if excitatory synaptic weights are restricted by normalization, the total excitatory current received by neurons within the memory assembly may be at the same level as the rest of the network. In that case, our vISP model would bring the activity of the memory assembly back to baseline level.

Network heterogeneity can also be promoted by heterogeneities in the structure of the network. We tested whether structural heterogeneity would be expressed in the neuronal activity across the network. Our vISP model resulted in a modulation of the overall network activity while maintaining the heterogeneity in the activity of the network. This diversity in network activity is in contrast with the observations from previous spike-timing dependent inhibitory synaptic plasticity models^15^. These models lead to a homogeneous neural network in which the underlying heterogeneity in the structure of the network is not exhibited in the network activity. Although this homogeneity could be remedied by assigning different targets to each neuron, the network would still not be flexible to adapt to, for example, different contexts. If target firing rates are fixed, spike-timing dependent models do not account for long-term rate-based encoding. This rigidity conflicts with experimental observations such as the changes in firing rate observed during perceptual learning^43;44^ or attention-dependent neuronal activity^45;46^. Although the firing rates are widely distributed when inhibitory connections follow our vISP model, the average ratio between excitatory and inhibitory currents converge to a unique level. Therefore, a voltage-dependent inhibitory synaptic plasticity rule does not set a target firing rate, but our simulations suggest that it might set a target E-I ratio.

A particularly interesting brain region in which to explore the consequences of neuronal diversity is the hippocampal CA1 sub-region. Pyramidal cells in this region have been reported to exhibit great variability from neuron to neuron in several features such as number of place fields, field sizes, and the proportion of the environment covered by the neuron^2^. Intracellular data suggests that neuronal excitability can account for the diversity in neuronal propensity of developing place fields and this propensity can determine the main behavioural features of these neurons^2^. Remarkably, CA1 pyramidal cells can be quickly and stably turned from silent to place cells^47^, inconsistently with a model that sets a unique target firing rate for each neuron, such as the sISP model. Moreover, inhibitory plasticity has been observed in the hippocampal network^29;48–50^. The wide range of neuronal features, however, is incompatible with a spike-based inhibitory synaptic plasticity model with a unique target firing rate. In our simulations, we observed that the combination of a heterogeneous distribution of pyramidal cell excitability with our vISP model can reproduce the experimentally observed variability in behavioural features. More importantly, our model supports changes in the network such as remapping caused by a change in environment. This diversity of features has been proposed as an important mechanism to ensure a combination of sparse coding — provided by the neurons with a low number of place fields — and a dense coding — provided by the neurons with a high number of place fields^1;2^. Furthermore, our vISP model supports the existence of a subset of cells that are active in one environment while silent in another. This might be essential for an efficient and easy context identification.

The model we propose aims to control the postsynaptic membrane voltage over a long timescale. Although there is experimental evidence that inhibitory plasticity depends on the postsynaptic membrane voltage, the exact form of this dependence has yet to be investigated. Further experiments would be necessary to constrain our model and further simulations could lead to more experimental predictions. The core conclusions from our simulations, however, depend solely on the fact that our inhibitory plasticity model does not depend only on spike timing.

The wider range of possible postsynaptic firing rates gives support for a more flexible network. While the E/I ratio and the maximum firing rates are forced onto the network, the different levels of postsynaptic firing rate allow for a more stable rate-based code. Since spike-based inhibitory plasticity models impose a fixed target firing rate, differences in firing rate can only be encoded in transient network dynamics. Following our vISP model, the network can converge to different states depending on the feedforward inputs. The neuronal response to sensory stimulation, for example, would abruptly increase at the stimulus onset but would decay to different levels depending on the amplitude of the stimulus. Therefore, the vISP model supports long-term, rate-based encoding. More importantly, our voltage-based model supports quick changes in rate-based codes caused, for example, by changes in context. Although a spike-based model could be combined with diversity in firing rates by assigning different target firing rates to each neuron, the entire network would still be constrained in terms of possible steady states. When changing environments, for example, hippocampal CA1 cells would fire, on average, at similar levels in both environments, regardless of the input received from Schaffer Collaterals or Entorhinal cortex connections.

Whereas our model imposes a natural maximum level of neuronal activity, it does not impose a minimum level. Consequently, pyramidal neurons are allowed to remain completely silent over long periods of time. Although it might be behaviorally relevant for some neurons to be silent^1;2^, biological circuits might have homeostatic mechanisms to ensure a minimum level of activity across the network. Recently, a local inhibitory synaptic plasticity rule has been proposed to control network-level activity^51^. Importantly, this input-dependent inhibitory synaptic plasticity (IDIP) does not impose a postsynaptic target firing rate and it may be combined with our voltage-dependent inhibitory plasticity model. Interestingly, this IDIP model can also be expressed as an intrinsic inhibitory plasticity model. A combination of an intrinsic plasticity model — following the IDIP model — and a voltage-dependent inhibitory synaptic plasticity model could regulate single neuron firing rate, allow for network diversity and adaptation, and modulate network-wide activity.

The interaction between excitatory and inhibitory synaptic plasticity has been shown to lead to complex dynamics^22^. Under the right conditions, this interaction leads to the development of receptive fields and co-tuning between excitation and inhibition^22^. Importantly, the details of the excitatory plasticity rule—more specifically, the choice of synaptic normalization—determine whether or not receptive fields are developed^22^. In all of our simulations, excitatory synaptic connections were fixed. It would be interesting to test whether the interaction between excitatory synaptic plasticity and our vISP model could lead to a more robust receptive field development. Since our inhibitory plasticity model supports different levels of postsynaptic firing rate, small differences in excitatory input would lead to small differences in firing rate that would not be compensated by a change in inhibitory synaptic weights. Therefore, this difference in excitatory synaptic weight would lead to a positive feedback loop and thus to the development of receptive fields.

The dynamics of the membrane voltage vary drastically depending on the location within the neuron. At the soma, action potentials follow a stereotypical shape spanning over a few of milliseconds^52^. At dendrites, however, NMDA spikes can last for dozens of milliseconds^52^. Therefore, our voltage-based inhibitory plasticity rule might have different effects depending on the targeting site of the inhibitory synapse. Since different types of interneurons generally target different layers of pyramidal cells, our vISP model could possibly explain the different learning rules observed for different types of interneurons^29–31^. Further simulations using physiologically detailed neurons would be required to confirm these stipulations.

Our voltage-based inhibitory plasticity model provides a mechanism to regulate network activity while supporting network diversity and flexibility. Our model imposes a natural maximum neuronal firing rate without setting a unique target value. More importantly, our inhibitory plasticity model allows for dynamic adaptation, an essential feature to account for context-dependent network states. Therefore, we provide a model that can be used to extend current computational approaches to simulate neural networks performing biologically-relevant tasks.

## Methods

### Neuron model

In our simulations, excitatory neurons are modelled by an adaptive exponential integrate-and-fire (AdEx) model^53^. As such, the neuronal membrane voltage *u* follows

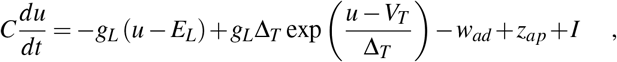

where *C* is the membrane capacitance, *g_L_* is the leak conductance, *E_L_* is the resting potential, Δ_*T*_ is the slope factor, *V_T_* is the threshold potential, and *I* is the total input current. The adaptation current *w_ad_* is described by

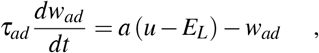

where *τ_ad_* is the adaptation time constant and *a* is a parameter. The depolarizing spike afterpotential *z_ap_* is set to *I_ap_* after a spike and decays exponentially with time constant *τ_z_* otherwise. The neuron spikes when its membrane voltage reaches a spiking threshold *V_th_*. At this point, the membrane voltage is reset to *V_reset_* and *w_ad_* is increased by an amount *b*. After spiking, the neuron’s membrane voltage is kept at *V_reset_* for a refractory time *τ_ref_*.

Additionally, we implement a conductance-based model for synaptic connections. Therefore, the total input current is described by

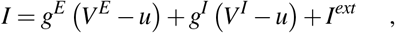

where *g^E^* is the excitatory synaptic conductance, *g^I^* is the inhibitory synaptic conductance, *V^E^* is the excitatory reversal potential, *V^I^* is the inhibitory reversal potential, and *I^ext^* is the external current. When the neuron receives an action potential from presynaptic neuron *j*, the postsynaptic conductance is increased by an amount 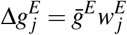, for excitatory synapses, and 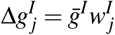, for inhibitory synapses. Both 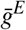 and 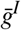 are parameters. The postsynaptic conductance decays otherwise with time constant *τ_E_*, for excitatory synapses, and *τ_I_*, for inhibitory synapses. The excitatory synaptic weights 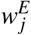 are fixed throughout the simulations and the inhibitory synaptic weights 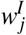 are updated following an inhibitory synaptic plasticity rule (see below).

This neuron model has been previously used in voltage-dependent excitatory plasticity models^24^ and has been shown to be important when used to fit experimental data^54^. All the parameters for the neuron model were adapted from previous studies^24;53^ and are given in table 1.

**Table 1:** Parameters summary 1. Neuron model, feedforward network, and synapse model.

| Neuron Model |  |  |
| --- | --- | --- |
| Name | Value | Description |
| $E_L$ | -70 mV | Resting potential |
| $g_L$ | 10 nS | Leak conductance |
| $\Delta_T$ | 4 mV | Slope factor |
| $V_T$ | -50 mV | Threshold potential |
| $\tau_{ad}$ | 150 ms | Adaptation time constant |
| $a$ | 2 nS | Subthreshold adaptation |
| $b$ | 80 pA | Spike-triggered adaptation |
| $V_{th}$ | -20 mV | Spiking threshold |
| $V_{reset}$ | -50 mV | Reset membrane potential |
| $\tau_{ref}$ | 5 ms | Refractory time |
| $C$ | 280 pF | Membrane capacitance |
| $C_I$ | 100 pF | Inhibitory membrane capacitance |
| $I_{ap}$ | 120 pA | Amplitude of afterpotential current |
| $\tau_z$ | 40 ms | Time constant of afterpotential current |
| Feedforward network parameters |  |  |
| Name | Value | Description |
| $N_E$ | 20 | Number of excitatory input neurons |
| $N_I$ | 20 | Number of inhibitory input neurons |
| Synapse Model |  |  |
| Name | Value | Description |
| $\tau_E$ | 5 ms | Decay constant for excitatory conductance |
| $\tau_I$ | 10 ms | Decay constant for inhibitory conductance |
| $\bar{g}_E$ | 1 nS | Maximum excitatory conductance amplitude |
| $\bar{g}_I$ | 1 nS | Maximum inhibitory conductance amplitude |
| $V^E$ | 0 mV | Excitatory reversal potential |
| $V^I$ | -80 mV | Inhibitory reversal potential |

Inhibitory neurons were modelled by a leaky integrate-and-fire (LIF) model. In this model, the membrane voltage *u* follows

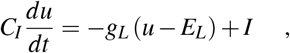

where *C_I_* is the inhibitory membrane capacitance.

### Synaptic plasticity model (vISP)

We propose a voltage-dependent inhibitory synaptic plasticity rule (vISP). Our synaptic plasticity model acts as a homeostatic mechanism to regulate the average postsynaptic membrane voltage. The weight of the synaptic connection from presynaptic inhibitory neuron *j* follows

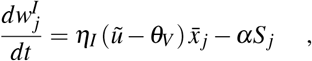

where *η_I_* is the inhibitory plasticity learning rate, *θ_V_* is a target membrane voltage, *α* is a parameter, and 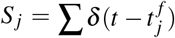 is the presynaptic spike train—where 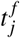 are the presynaptic spiking times. The presynaptic trace 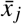 is increased by 1 whenever there is a presynaptic spike and decays exponentially otherwise with time constant *τ_trace_*. The variable 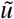 is an exponential low-pass-filtered version of the postsynaptic membrane potential

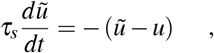

where *τ_s_* is a slow time constant.

### Synaptic plasticity model (sISP)

To compare the effect of our voltage-based inhibitory plasticity model with previous models, we run some of the simulations under the same conditions but replacing the plasticity rule with an spike-timing-dependent inhibitory synaptic plasticity model (sISP)^15^. Under this model, the weight of the synaptic connection from presynaptic inhibitory neuron *j* to postsynaptic neuron *i* is updated such that

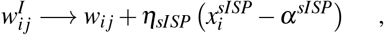

for every presynaptic spike, and

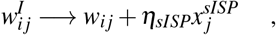

for every postsynaptic spike, where *η_sISP_* is the sISP learning rate, *α^sISP^* is the depression factor, and *x^sISP^* is the synaptic trace which is increased by 1 whenever the neuron spikes and decays exponentially otherwise with time constant *τ_sISP_*.

### Recurrent networks

We simulate a network of *N_E_* excitatory and *N_I_* inhibitory neurons receiving inputs from *N_pre_* excitatory neurons. The neurons in the recurrent network are simulated as AdEx and LIF neurons as described above. Input neurons are modelled as Poisson neurons with a constant firing rate *λ_pre_*. These neurons are connected to neurons in the recurrent network — both excitatory and inhibitory neurons — with probability *p_pre_*. Within the recurrent network, neurons are connected with probability *p_EE_* (E-E connections), *p_EI_* (I to E connections), *p_IE_* (e to I connections), and *p_II_* (I-I connections).

#### Figure 3 - randomly connected network

Initial excitatory to excitatory synaptic weights are drawn from a log-normal distribution with parameters *μ* = 0 and *σ* = 0.5 (mean and standard deviation of the associated normal distribution, respectively). Initial inhibitory synaptic weights onto excitatory cells are drawn from a log-normal distribution with parameters *μ* = 0 and *σ* = 1. Excitatory to inhibitory synaptic weights are set to 3 whilst inhibitory to inhibitory synaptic weights are set to 30. The simulation is split into three main stages. During the first stage (random network), the network is simulated for 100 seconds. At the end of this stage, a subset of 200 neurons were selected and the connections amongst these neurons were multiplied by 3. During the second stage of the simulation (memory formation), the network was simulated for a further 100 seconds. During the third stage (memory retrieval), half of the neurons in the memory assembly formed in the previous stage were overstimulated. This extra stimulation is implemented by simulating an additional 0.4*N_pre_* input neurons with a firing rate 10 times higher than the other input neurons. These neurons are then transiently connected to the selected neurons within the memory assembly. This third stage is simulated for 12.5 seconds. We then simulate the network without this extra input for another 12.5 seconds. Inhibitory plasticity is active throughout the entire simulation.

#### Figure 4 - heterogeneously connected network

In these simulations, excitatory neurons in the recurrent network were heterogeneously connected. The probability that neuron *i* was connected to neuron *j* was given by 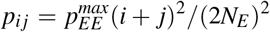, where 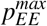 is the maximum probability of connection between two neurons. Due to this connection probability, the first neurons in the network are sparsely connected to the rest of the network whereas the last neurons are densely connected (figure 4B). All the excitatory synaptic weights were then set to 3. The remaining weights were defined as in figure 3.

### Hippocampal network

To simulate the CA1 hippocampal network, we simulated a network of excitatory neurons connected to inhibitory neurons. Excitatory neurons are not connected to other excitatory neurons whereas the other connections (E-I, I-E, and I-I) are simulated in the same way as in the recurrent network simulations. All neurons in the CA1 network receive inputs from Poisson neurons representing CA3 pyramidal cells. The firing rates of these CA3 cells are modulated by the position of the animal along a linear track following a Gaussian shape with amplitude *λ_E_* and width *σ*_*CA*3_ = 1. The total length of the track is *L* = 100 and the CA3 place fields are evenly distributed along the track. CA3 neurons are then randomly connected to CA1 neurons with the same probability as in figure 3 and 4. For each CA1 pyramidal cell, we set its threshold potential *V_T_* to a value drawn from a uniform distribution from −50 mV to −20 mV. This value is then kept fixed throughout the entire simulation. When simulating environment changes, we redraw both the connections and the synaptic weights from the CA3 to the CA1 network. We use the same probability distributions in both environments A and B.

### Parameters and simulations

All data and software supporting the findings of this study are available on ModelDB. The parameters for all the simulations can be found in tables 1-2.

**Table 2:** Parameters summary 2. Plasticity models and recurrent network.

| Plasticity model (vISP) |  |  |
| --- | --- | --- |
| Name | Value | Description |
| $\eta_I$ | $1 \times 10^{-3} \text{ ms}^{-1} \text{ mV}^{-1}$ | Inhibitory plasticity learning rate |
| $\theta_V$ | -65 mV | Target membrane voltage |
| $\alpha$ | $2 \times 10^{-4}$ | Depression parameter |
| $\tau_{trace}$ | 5 ms | Decay time constant for presynaptic trace |
| $\tau_s$ | 200 ms | Slow decaying time constant |
| Plasticity model (sISP) |  |  |
| Name | Value | Description |
| $\eta_{sISP}$ | 0.3 | Inhibitory plasticity learning rate (sISP model) |
| $\alpha^{sISP}$ | 0.08 | Presynaptic offset |
| $\tau_{sISP}$ | 20 ms | Decay constant for synaptic trace |
| Recurrent network parameters |  |  |
| Name | Value | Description |
| $N_{pre}$ | 1000 | Number of excitatory input neurons |
| $N_E$ | 1000 | Number of excitatory neurons in the recurrent network |
| $N_I$ | 250 | Number of inhibitory neurons in the recurrent network |
| $\lambda_{pre}$ | 5 Hz | Firing rate of input neurons |
| $p_{pre}$ | 0.2 | Probability of connections from input neuron |
| $p_{EE}$ | 0.4 (Fig3); 0.0 (Fig5-6) | Probability of E-E connections (except Fig 4) |
| $p_{EE}^{max}$ | 0.9 (Fig4) | Maximum probability of E to E connections for figure 4 |
| $p_{EI}$ | 0.2 | Probability of I to E connections |
| $p_{IE}$ | 0.2 | Probability of E to I connections |
| $p_{II}$ | 0.2 | Probability of I-I connections |
| $w_{pre}^E$ | 4 | synaptic weights from input to excitatory neurons |
| $w_{pre}^I$ | 1 | synaptic weights from input to inhibitory neurons |

## Notes

### Competing Interest Statement

The authors have declared no competing interest.

