## Supplementary Material for "Voltage-based inhibitory synaptic plasticity: network regulation, diversity, and flexibility"

### 1 Figures

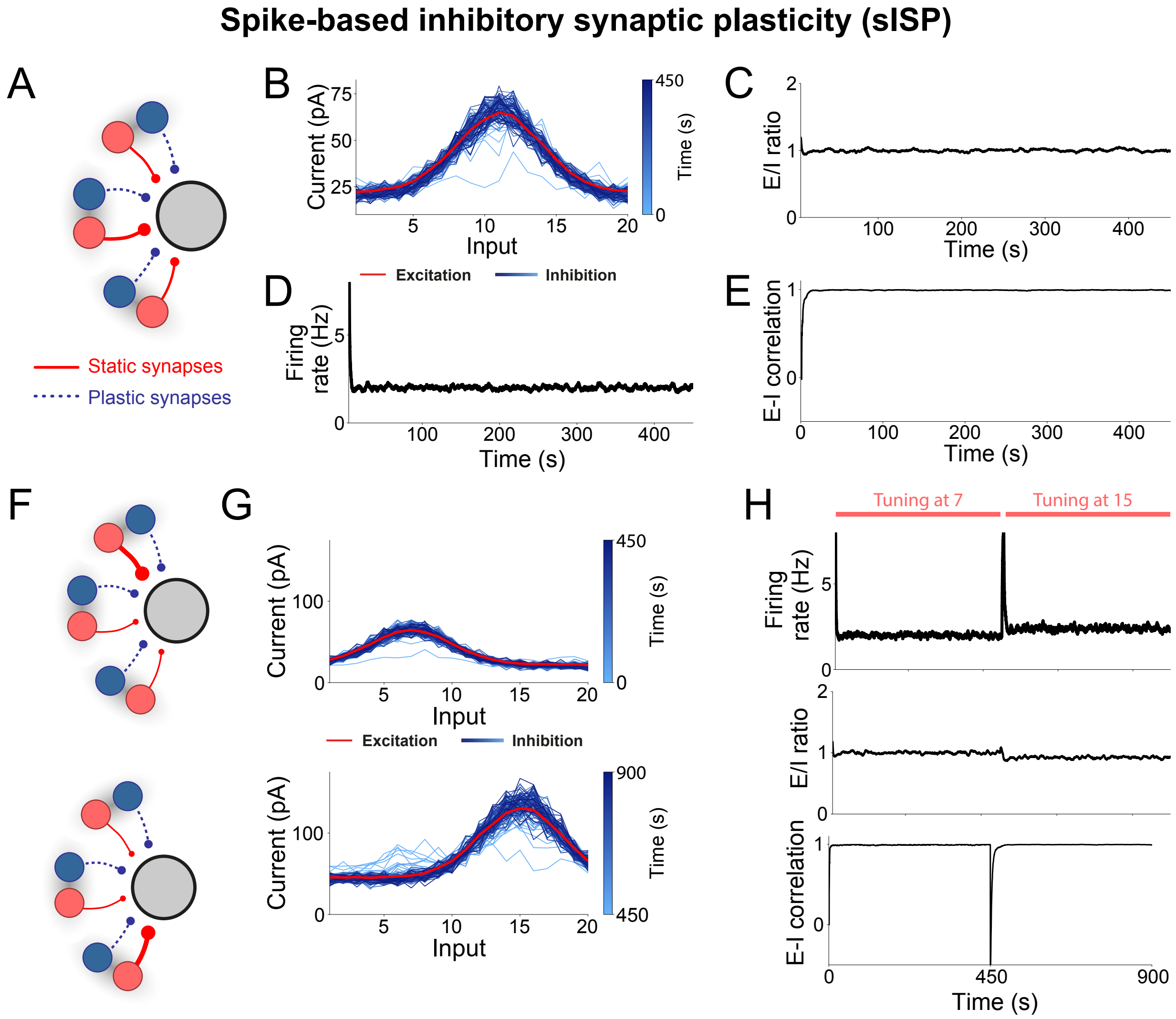


**Supplementary Figure 1**: **(Related to figure 2) sISP and correlated E-I inputs lead to co-tuned excitatory and inhibitory receptive fields.** **(A)** Network diagram. We simulate a feedforward network composed of a set of excitatory (red) and inhibitory (blue) neurons projecting onto one postsynaptic excitatory cell (grey). Each excitatory neuron is associated with an inhibitory counterpart such that they follow the same time-varying firing rate. The firing rates of different pairs of input neurons vary over time but are kept, on average, at the same level. Inhibitory connections follow a spike-based inhibitory synaptic plasticity (sISP) model. **(B)** Evolution of inhibitory receptive field. Synaptic weights for each inhibitory (blue) and excitatory (red) input neuron. The different shades of blue represent snapshots of the inhibitory weights at different times throughout the simulation. **(C)** Ratio between excitatory and inhibitory currents onto the postsynaptic cell as a function of time. The curve is smoothed using a rolling average over 100 ms. **(D)** Postsynaptic firing rate as a function of time. **(E)** Correlation between excitatory and inhibitory receptive fields as a function of time.

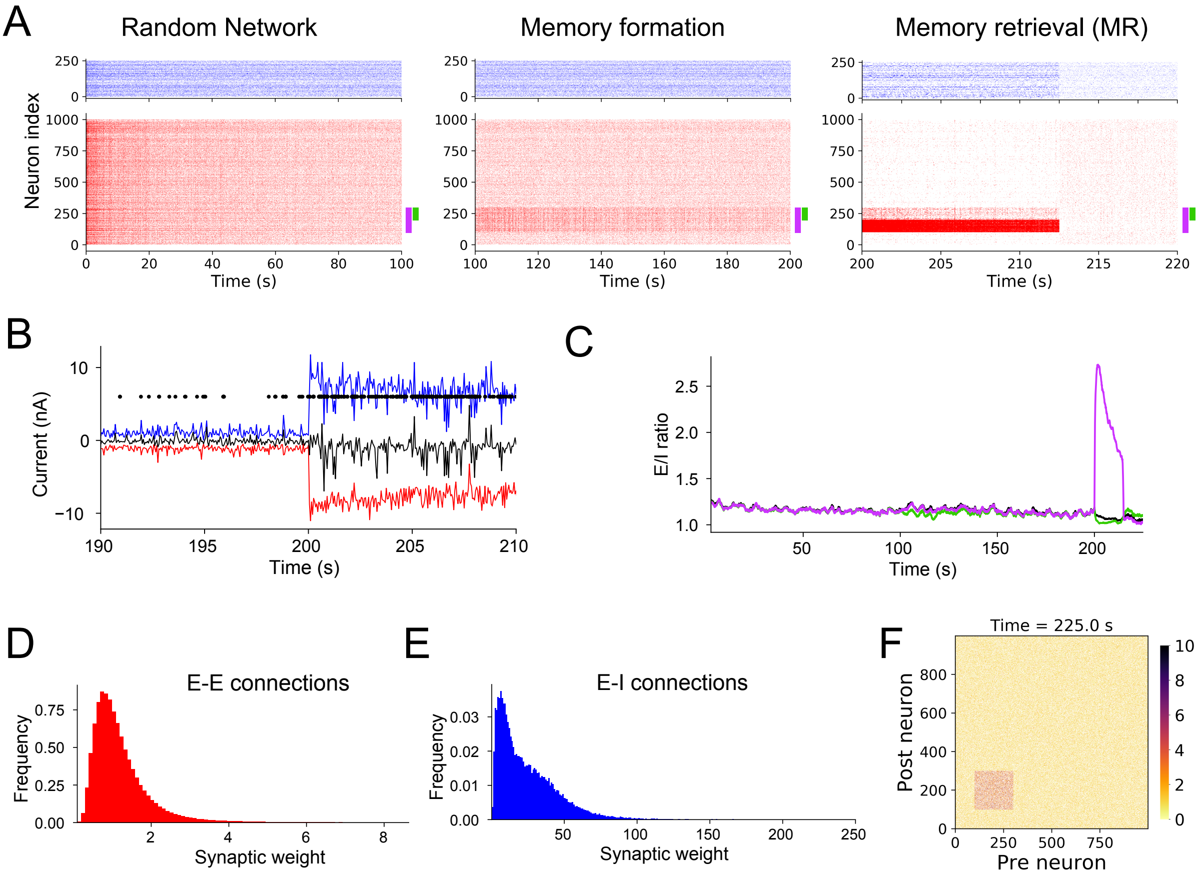


**Supplementary Figure 2**: **(Related to figure 3) . (A)** Inhibitory (blue) and excitatory (red) spikes for the first stage of the simulation (left), the stage following the memory imprinting (middle), and the memory retrieval stage (right) of the simulation in figure 3. Purple: memory assembly whose neurons are strongly connected amongst themselves after $t = 100$ s. Green: half of the neurons in the memory assembly that does not receive an extra external input at $t = 200$ s. **(B)** Inhibitory (blue), excitatory (red), and net (black) currents received by a random neuron within the second half of the memory assembly (green bar in A). Black dots indicate a spike. The currents shown are the same as in figure 3C zoomed in around the memory retrieval. **(C)** Average ratio between excitatory and inhibitory currents for the neurons in the memory assembly (purple), the non-stimulated neurons in the assembly (green), and the neurons outside of the assembly (black). **(D)** Distribution of excitatory to excitatory synaptic weights. **(E)** Final distribution of inhibitory to excitatory weights. **(F)** Excitatory weight matrix at $t = 225$ s (end of simulation).

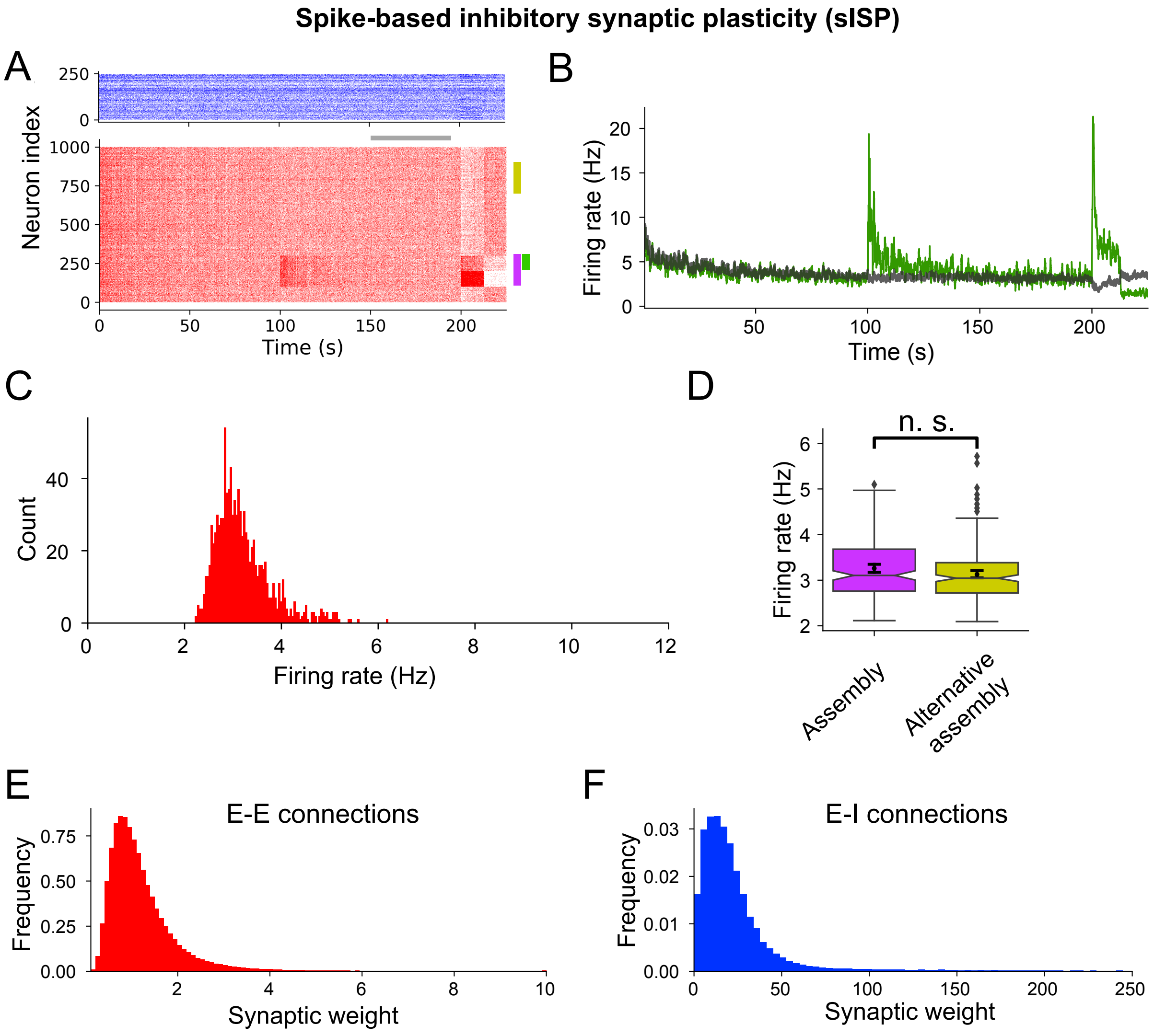


**Supplementary Figure 3**: **(Related to figure 3) The activity of memory assemblies fade back to background level under sISP.** **(A)** Inhibitory (blue) and excitatory (red) spikes for one example simulation. The memory formation stage starts at 100 seconds and the extra input (memory retrieval) is applied at 200 seconds for 5 seconds. Purple: memory assembly whose neurons are strongly connected amongst themselves after $t = 100$ s. Green: half of the neurons in the memory assembly that does not receive an extra external input at $t = 200$ s. Yellow: an alternative set of cells of the same size as the memory assembly. The grey bar indicates the interval over which the average firing rates are measured in C and D. **(B)** Firing rate for the second half of the neurons in the memory assembly (green bar in A) and background activity for the neurons outside of the memory assembly. **(C)** Distribution of firing rates measured between 150 and 190 seconds (grey bar in A) across all excitatory neurons. **(D)** Comparison between the average firing rate of the memory assembly (purple bar in A) and an alternative set of cells (yellow bar in A). The average firing rate is measured from 150 to 190 seconds. The firing rates are not significantly different after network stabilization ($p = 0.0538$, Kruskal-Wallis H-test, $n = 200$ neurons). **(E)** Distribution of excitatory to excitatory synaptic weights. **(F)** Final distribution of inhibitory to excitatory weights.

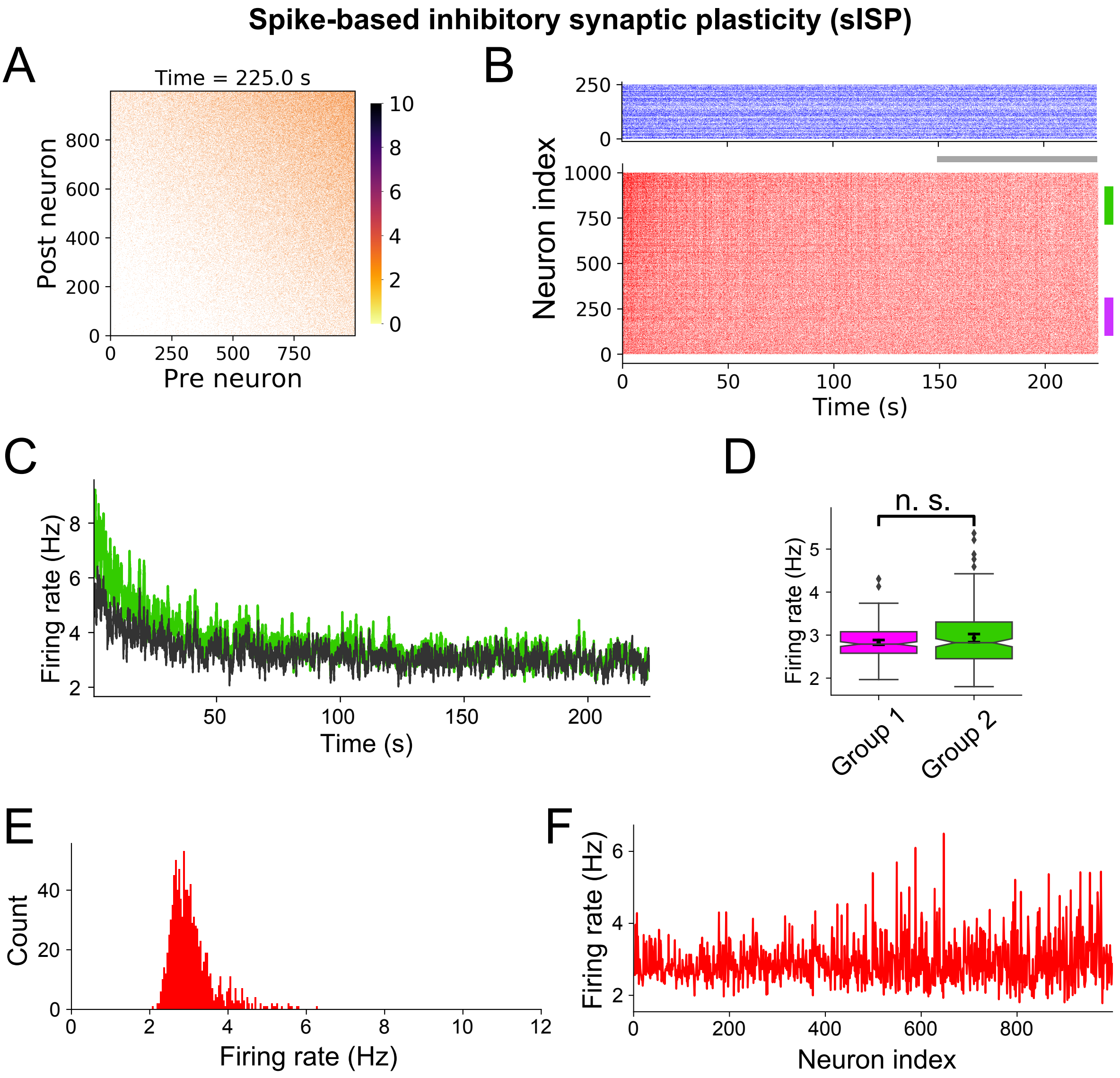


**Supplementary Figure 4**: **(Related to figure 4) sISP suppresses diversity in a heterogeneous recurrent network.** **(A)** Excitatory synaptic connectivity matrix. All the connections have the same strength, but the neurons are not uniformly connected. Neurons are sorted by their propensity of forming connections with other neurons. **(B)** Inhibitory (blue) and excitatory (red) spikes. Purple: set of 200 neurons with low propensity of forming connections with other neurons in the network. Green: set of 200 neurons with high propensity of forming connections with other neurons in the network. The grey bar indicates the interval over which the average firing rates are measured in D, E and F. **(C)** Firing rate for a set of highly connected neurons (green bar in B) and background activity measured across all the other neurons. **(D)** Comparison between the average firing rate of a set of weakly connected neurons (group 1, purple bar in B) and a set of highly connected neurons (group 2, green bar in B). The average firing rate is measured from 150 to 220 seconds. The firing rates are not significantly different after network stabilization ($p = 0.466$, Kruskal-Wallis H-test, $n = 200$ neurons). **(D)** Distribution of firing rates measured from 150 and 220 seconds (grey bar in B) across all excitatory neurons. **(E)** Firing rates shown in E for each neuron. Neurons are sorted by their propensity of forming connections with other neurons.

#### Spike-based inhibitory synaptic plasticity (sISP)

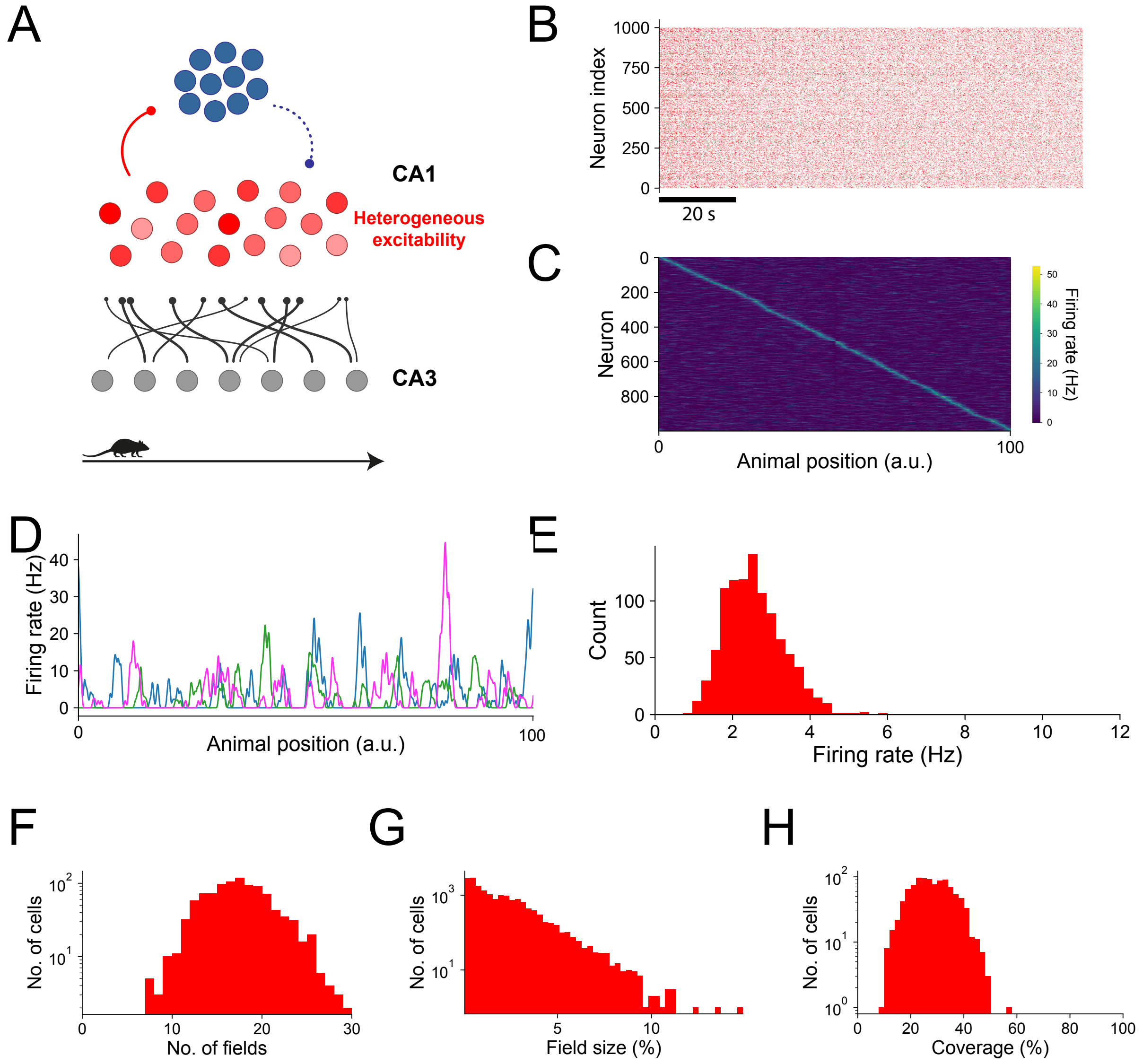


**Supplementary Figure 5**: **(Related to figure 5) sISP constrains CA1 place field diversity.** **(A)** Network diagram. CA1 pyramidal cells (red) receive inputs from CA3 excitatory cells (grey) and are recurrently connected via inhibition (blue). Each excitatory cell is assigned a level of excitability such that their spiking thresholds are drawn from a uniform distribution. CA3 place fields are uniformly distributed and spam the entire linear track. Connections from CA3 to CA1 pyramidal neurons and from pyramidal neurons to interneurons are random and fixed. The inhibitory connections onto CA1 pyramidal cells are initialised randomly and follow a spike-dependent inhibitory synaptic plasticity rule. **(B)** CA1 pyramidal cell spikes for the entire simulation. The animal travels across the entire linear track in 10 s and is placed back at the initial position instantly. **(C)** CA1 pyramidal cell activity as a function of the animal position. Neurons were sorted by the position of their maximum firing rate. The activity is an average over the final 5 laps of simulation. CA1 place fields span over the entire linear track. **(D)** Example place fields for 3 CA1 pyramidal cells (each color represents one cell). Each cell can have multiple place fields across the track and their amplitudes can vary significantly. **(E)** Distribution of average firing rates across CA1 pyramidal cells. Firing rates were measured over the final 5 laps of simulation. The sISP model constrains network activity to a narrow range of values. **(F)** Distribution of number of fields. **(G)** Distribution of field size compared to the total track. **(H)** Distribution of the total coverage of each neuron compared to the total track.

#### Spike-based inhibitory synaptic plasticity (sISP)

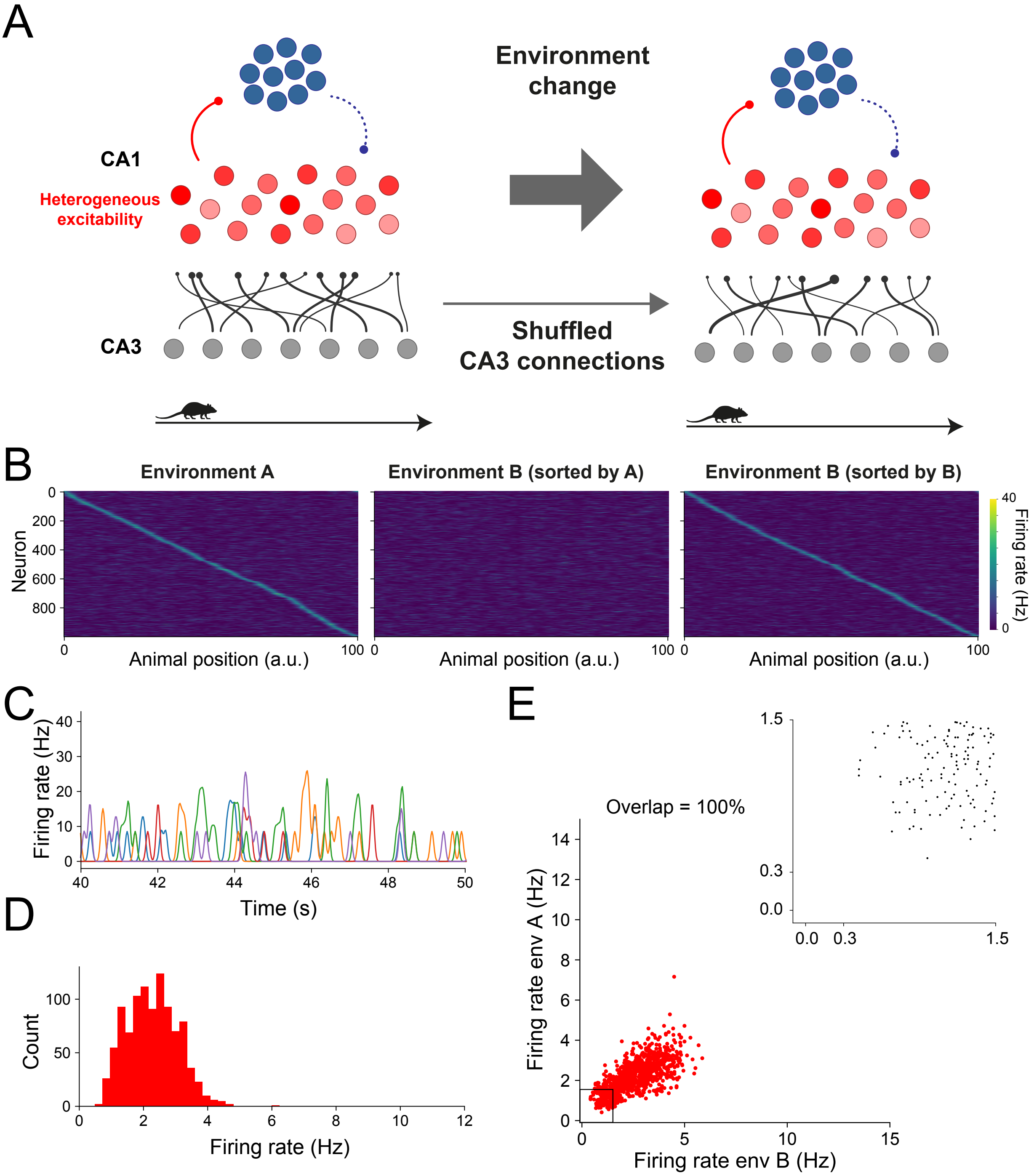

**Supplementary Figure 6**: **(Related to figure 6) sISP does not support flexible neuronal remapping. (A)** Network diagram and simulation protocol. The network is simulated as in figure 5. After half of the simulation time, the connections from CA3 to CA1 pyramidal cells are instantly re-drawn from the same distribution as before to simulate an environment change. The network is then simulated as in the first half of the simulation. All the other parameters are kept constant throughout the entire simulation, including the excitability of CA1 pyramidal cells. Inhibitory connections follow a spike-based inhibitory synaptic plasticity (sISP) model. **(B)** CA1 pyramidal cell activity as a function of the animal position for the first half (environment A) and the second half (environment B) of the simulation. Neurons were sorted by the position of their peak in either environment A (left and middle panels) or environment B (right panel). The activity shown is an average over the final 5 laps of simulation in each environment. **(C)** Example place fields for 10 CA1 pyramidal cells in environment B. Similarly to before the change in CA3 connections, each cell can have multiple place fields across the track and their amplitudes can vary significantly. **(D)** Distribution of average firing rates across CA1 pyramidal cells in environment B. Firing rates were measured over the final 5 laps of simulation. **(E)** Firing rate in environment A versus firing rate in environment B for all the CA1 pyramidal cells. The overlap represents the proportion of cells that either active or silent in both environments (i.e. the proportion of cells that do not change their status as silent or place cells). Therefore, none of the cells are active in only one of the environments. Inset: firing rate within the range between zero and 1.5 Hz. Neurons with an average firing rates below 0.3 Hz were considered silent. All the neurons were active in both environments.
